# Detection of rare plasmid hosts using a targeted Hi-C approach

**DOI:** 10.1101/2023.11.30.569469

**Authors:** Salvador Castaneda-Barba, Benjamin J. Ridenhour, Eva M. Top, Thibault Stalder

## Abstract

Despite the significant role plasmids play in microbial evolution, there is limited knowledge of their ecology, evolution, and transfer in microbial communities. Therefore, we developed and implemented a novel approach to identify rare plasmid hosts by combining Hi-C, a proximity ligation method, with enrichment for plasmid-specific DNA. We hereafter refer to this Hi-C enrichment approach as Hi-C+. Our experimental design mimicked scenarios in which the transfer of an antimicrobial resistance plasmid from a donor to a recipient in soil was increasingly rare. We established that Hi-C can link a plasmid to its host in soil when the relative abundance of that plasmid-host pair is as low as 0.001%. The Hi-C+ method further improved the detection limit of Hi-C 100-fold and allowed identification of plasmid hosts at the genus level. Therefore, Hi-C+ will facilitate the exploration of the ecological and evolutionary pathways that affect the spread of plasmids in natural environments.

**Teaser:** In this study we demonstrate that a target-enriched Hi-C approach can identify rare hosts of a given plasmid in soil.

## Introduction

Plasmids are mobile genetic elements that replicate separately from the bacterial chromosome and can transfer by conjugation to other closely and distantly related bacteria (*1*, *2*). It is now evident that horizontal gene transfer mediated by plasmids plays a crucial role in enabling bacteria to rapidly adapt to changing environments (*3*, *4*). An important example is the role of plasmids in spreading antibiotic resistance. Antibiotic-resistant bacteria already inflict a significant health burden, having caused an estimated 1.27 million deaths worldwide in 2019 (*5*). Plasmids can carry genes that encode resistance to 10 or more antibiotics (*6*), including so-called “drugs of last resort” (*7–10*). Recently, these multi-drug-resistance (MDR) plasmids have been increasingly involved in treatment failures (*11*), emphasizing the importance of understanding the conditions that facilitate their mobilization to human pathogens.

To understand the spread of antimicrobial resistance and the horizontal transfer of any trait between bacteria, it is critical not to limit the study of plasmid transmission to human pathogens alone. Antimicrobial resistance is a global issue that goes beyond human health as many antibiotic resistance genes of concern in human pathogens are also prevalent in bacteria from animal and environmental habitats (*12*). From these settings, plasmids facilitate the spread of antibiotic resistance genes across habitat boundaries to human pathogens (*13–15*). Therefore, it is crucial that we determine the conditions that facilitate the spread and persistence of plasmids in their natural habitats (*16*). One critical challenge is to identify the bacteria that can acquire and retain MDR plasmids in the environment, as they can facilitate both the long-term persistence of MDR plasmids and their spread to other members of the microbiome (*17–19*). Unfortunately, current methods are limited in their ability to identify plasmid hosts *in situ*, leaving an important knowledge gap in the field of plasmid ecology.

One important goal is to be able to identify the bacterial hosts that have acquired a focal MDR plasmid in an environment such as soil. Traditionally, one can isolate and identify new plasmid recipients, called transconjugants (*20–22*), through plating or enrichment of bacteria that express the phenotype of interest conferred by a focal plasmid. However, this technique is limited to the culturable fraction of bacteria, which is known to be less than 1% in soil (*23*). Several groups have successfully marked their focal plasmid with a gene encoding a fluorescent protein, followed by flow cytometry to track and even identify putative new transconjugants (*24–27*). Modifying the plasmid this way has serious drawbacks: (i) it may change the transferability and persistence of the plasmid due to the inserted sequence (*28*), and (ii) fluorescence of these reporters can be strain and environment specific, making detection in communities difficult and biased (*29*, *30*). Moreover, given that the plasmid has been genetically modified, monitoring it in the field or a human microbiome may not be permitted. One could also quantify the presence of the focal plasmid using qPCR (*31–33*), or use shotgun sequencing to assess the relative abundance of the plasmid, as is frequently done for antibiotic resistance genes (*34–36*). Unfortunately, neither of these methods would give us information on the identity of the bacteria that have acquired the plasmid. Altogether, these methods are limited in their ability to identify the hosts of a plasmid in microbial communities, thereby hampering us from tracking the plasmid spread.

The development of chromosomal conformation capture (or proximity ligation) approaches for microbiome exploration, such as Hi-C and meta3C, has opened the door to further our insight into plasmid-mediated resistance spread in natural habitats. These approaches rely on crosslinking DNA within the cell, prior to cell lysis (Fig. 1). The physical contacts produced by the crosslinks provide quantitative and objective information on whether two segments of DNA share the same cellular compartment, without requiring any prior knowledge of the bacterial community to which the method is applied (*37–39*). This information can help understand plasmid and phage-host interactions and aid in reconstructing the reservoirs of antibiotic resistance genes and plasmids in wastewater as well as in human and pig gut microbiomes (*40–46*).

**Fig. 1:**
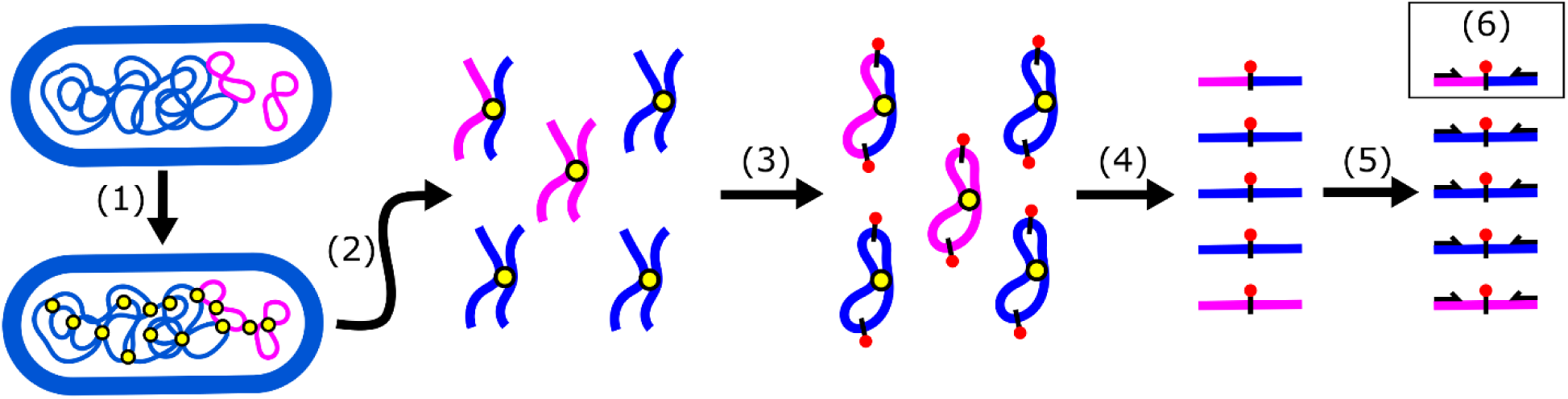
Hi-C method. 1) Before cell lysis, DNA is crosslinked with formaldehyde. Cross-links between DNA in proximity are depicted as yellow circles; (2) cells are lysed, DNA is then purified and cut with a restriction enzyme; (3) cut restriction sites are filled in using biotin-labeled nucleotides (red circles) and are then religated; (4) DNA is sheared, and biotin-labeled fragments are enriched; (5) sequencing adapters are added, and Hi-C fragments are sequenced. The paired-end Hi-C reads are depicted as black arrows. (6) Paired-end reads from chimeric fragments of DNA that linked the plasmid to its host can be used to identify that the pink plasmid was present in the blue bacteria.

Despite its implementation for microbiome analysis, an important limitation of Hi-C is its sensitivity to detect specific targets in a microbial community. This is particularly important for monitoring plasmid spread, as transfer in natural habitats occurs at very low frequencies (*47–49*). For example, the transfer frequency of conjugative plasmid pJP4 from a known donor to recipient in nonsterile soil was 10^-6^ (*48*). Furthermore, many resistance plasmids within microbial communities are present in members that are in low abundance (*17*, *19*). This presents a challenge for researchers interested in identifying ecological and evolutionary trajectories of a focal plasmid in a natural community. To overcome this challenge, we paired Hi-C with a target enrichment approach, hereafter referred to as Hi-C+. We first determined the detection limit of Hi-C for linking a plasmid to its host. Subsequently, we showed that Hi-C+ can (i) improve the detection limit of Hi-C by enriching the Hi-C DNA library with plasmid-specific DNA, and (ii) facilitate genus-level identification of new plasmid hosts.

## Results

### Hi-C detects a plasmid-host pair present at a relative abundance of 10^-5^

To determine the detection limit of Hi-C, we designed an experiment meant to mimic a scenario where a plasmid-carrying bacterial strain is introduced into soil, and subsequently transfers its plasmid to a soil bacterium (Fig. 2, Table 1). We refer to the initial plasmid-carrying strain as the donor, the bacteria that can acquire the plasmid as the recipients, and those to which the plasmid has transferred as the transconjugants. Rather than inoculating donors and recipients and letting plasmid transfer occur at unknown frequencies, we opted to inoculate donors, recipients, and transconjugants at a series of known densities (Fig. 2B, Table 1). This allowed us to accurately assess the Hi-C method and determine its limit in detecting contacts between DNA from a known plasmid and the chromosomes of the transconjugants and donors.

**Fig. 2:**
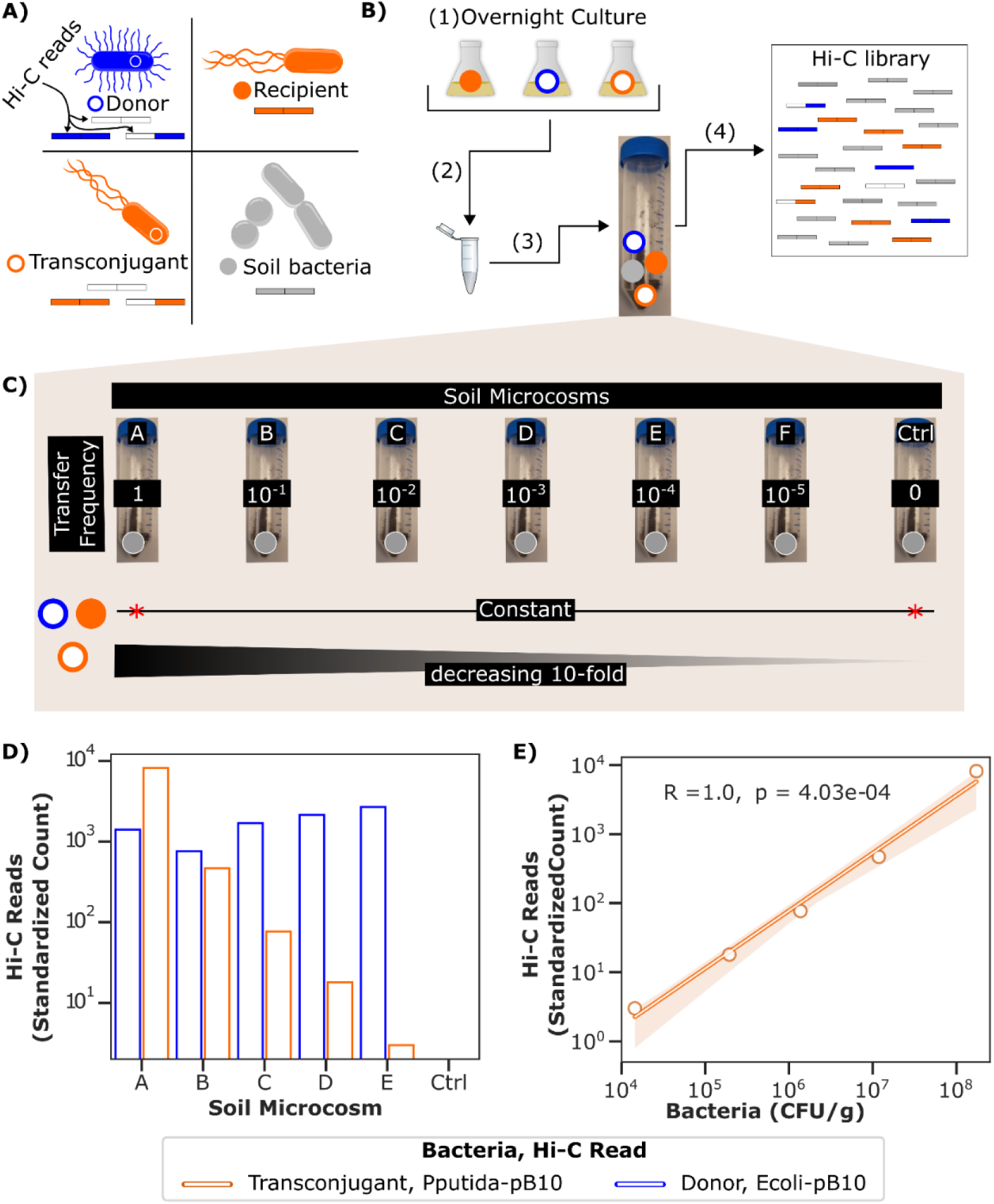
Overview of soil microcosm setup and detection of transconjugants and donors by Hi-C. A) Legend for colors used to represent the donor, recipient, transconjugant, and soil bacteria. The types of Hi-C reads that can be generated by each bacterium are also depicted. B) To accurately quantify the limits of Hi-C in detecting transconjugants, we set up soil microcosms to which we added a specific amount of donor, recipient, and transconjugant. An experimental flow chart of the soil microcosms is depicted: 1) Donor, transconjugant, and recipient were grown overnight. 2) Appropriate amount of each bacterium was pooled and 3) subsequently added to soil. A visual representation of the composition of each microcosm is depicted in panel C. 4) To limit opportunities for plasmid transfer in our experimental design, soil from each microcosm was immediately crossed-linked and underwent the Hi-C library preparation. Hi-C libraries were then sequenced. C) Visualization of soil microcosm composition. The decreasing transconjugant frequency simulates scenarios where plasmid transfer is increasingly rare and allows us to measure the detection limit of Hi-C. *No recipients were added to soil microcosm A, Ctrl soil microcosm contained Recipient, a plasmid-free Donor strain and no Transconjugant. D) Reads linking the plasmid to the transconjugant and donor chromosome were detected in all soil microcosms tested, except for the control. To account for differences in sequencing depth, Hi-C read count was standardized to the lowest sequencing depth. E) Relationship between transconjugant bacteria plated (Table 1) and Hi-C reads observed. A Pearson correlation coefficient was computed and is displayed in the top left.

**Table 1:**
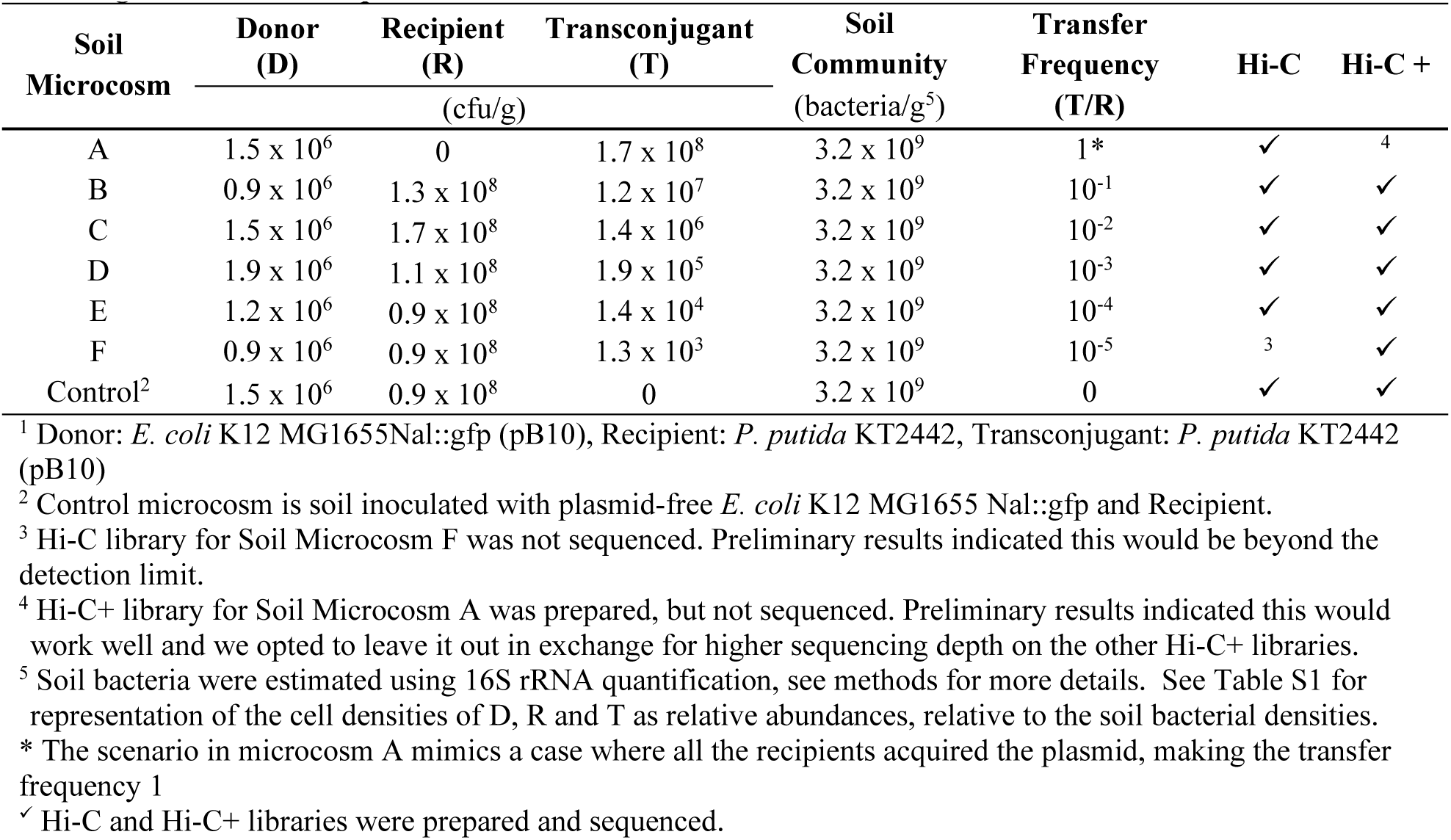
Soil microcosms imitating plasmid transfer scenarios with densities of each bacterial strain^1^ added and targeted ‘transfer frequencies’.

We set up seven soil microcosms with various mock plasmid transfer scenarios (Table 1, Fig. 2B, C). Each microcosm represents a scenario where the plasmid has transferred to an increasingly smaller fraction of the recipients, resulting in a declining transfer frequency and overall proportion of transconjugants. As a focal plasmid in this setup, we used MDR IncP plasmid pB10::rfp, hereby referred to as pB10. As recipients, we used a *Pseudomonas putida* KT2442. As plasmid donors and transconjugants, we respectively used *Escherichia coli* K-12 MG1655Nal::gfp (pB10) and *P. putida* KT2442 (pB10). An overview of the experimental design for our soil microcosms is shown in Fig. 2. A-C.

Our first goal was to determine if the Hi-C method could detect Hi-C reads associating pB10 with the decreasing transconjugant and constant donor chromosomes in each of the soil microcosms. To do this, we generated and sequenced Hi-C libraries from soil microcosm A through E and control (Table 1). When analyzing the paired-end Hi-C reads, we specifically looked for those where one aligned perfectly to the *P. putida* KT2442 or *E. coli* MG1655 reference sequences and the other to pB10. These reads indicate the presence of the plasmid in the transconjugant and donor bacteria, and are respectively denoted as Pputida-pB10 and Ecoli-pB10 Hi-C reads. This hyphenated nomenclature is used hereafter and is intended to indicate the two genomes to which paired-end reads aligned. We detected Pputida-pB10 and Ecoli-pB10 Hi-C reads in all but the control Hi-C libraries (Fig. 2D). Importantly, transconjugants were still detected even when the donor was 100 times more abundant (E in Table 1 and Fig. 2). Additionally, there was a strong correlation between the amount of transconjugants added and observed Pputida-pB10 Hi-C reads (Fig. 2E). Our results demonstrate that Hi-C can distinguish a plasmid present in multiple hosts in a natural soil community, even when one host is present at a low relative abundance.

Ideally, to monitor the spread of antimicrobial resistance, researchers should be able to enumerate new plasmid-host pairs in soil or other environments and compare observations between treatments or over time. To address this, we built a quantitative model that relates Hi-C reads to bacterial counts, using the data collected from the sequenced soil microcosms in conjunction with plate count data (Fig. 2D and Table 1). To account for the effect of sequencing depth, we first randomly subsampled the paired-end reads from each soil microcosm at the following sequencing depths; 50, 60, 70, 80, and 90 million paired-end reads. We then performed negative binomial regression (Equation 1) with a response variable of bacterial count (cfu/g) and predictor variables Pputida-pB10 Hi-C reads (*z-score* = 16891, *p* < .001) and sequencing depth (*z-score* = -3092, *p* <. 001).

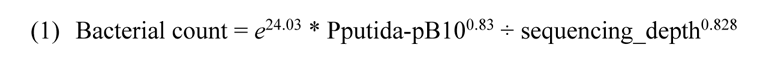

Using this model will enable future estimations of the prevalence of an introduced plasmid in an indigenous bacterial community.

Perhaps the most pressing question prior to using Hi-C to detect *in situ* plasmid transfer is whether a plasmid-host pair is present at a high enough relative abundance in the community to be detected in a sequenced library. Therefore we determined the detection limit of Hi-C by calculating the minimum number of plasmid hosts needed in a bacterial community to allow detection of at least one Hi-C read that links the focal plasmid to its host with high confidence. First, using Poisson probability we identified that a library needed to have a mean of 9 plasmid-host DNA fragments, for there to be a high probability (P(X>0) = 99.99%) of detecting at least one plasmid-host Hi-C read within a sequencing run. Using equation 1, we calculated that given a sequencing depth of 90 million paired-end reads, this corresponds to 3.3 x 10^4^ bacteria/g of soil. Presented as a relative abundance of the number of bacteria in our soil microcosm, 3.2 x 10^9^ bacteria/g of soil, this equates to 10^-5^. In the context of our experimental setup, this detection limit would be between microcosms D and E. The detection of transconjugant Hi-C reads in scenario E shows that reads can still be detected below the identified limit, but this observation has a lower probability of occurring. It is also important to note that this detection limit is dependent on the desired number of observed Hi-C reads. Maintaining the same sequencing depth but increasing the desired observed Hi-C reads to 10 or 100 would respectively result in detection limits of 1.1 x 10^5^ and 7.1 x 10^5^ bacteria/g of soil. Altogether, our results demonstrate that when a plasmid-host pair is present in a community at a relative abundance of 10^-5^, we can expect to detect it in 90 million paired-end reads.

### Hi-C+, enriching a Hi-C library with plasmid DNA through target capture

In natural microbiomes, plasmid transfer is often rare. Transconjugant proportions can be below 10^-5^ (*47–49*), the detection limit of Hi-C (Fig. 2). To circumvent this limitation, we set out to improve the detection limit of this method for studies that monitor one focal plasmid in a bacterial community. This was accomplished by combining Hi-C with a target enrichment approach, a method we refer to as Hi-C+ (Fig. 3). Hi-C+ consisted of an in-solution hybridization capture of target DNA using biotinylated custom RNA baits that are complementary to the entire 68-kb sequence of our focal plasmid, pB10. It was applied directly to DNA from the Hi-C libraries of soil microcosms B through F and the control microcosm, yielding so-called Hi-C+ libraries (see Table 1). To minimize experimental variation, all reactions were carried out in parallel.

**Fig. 3:**
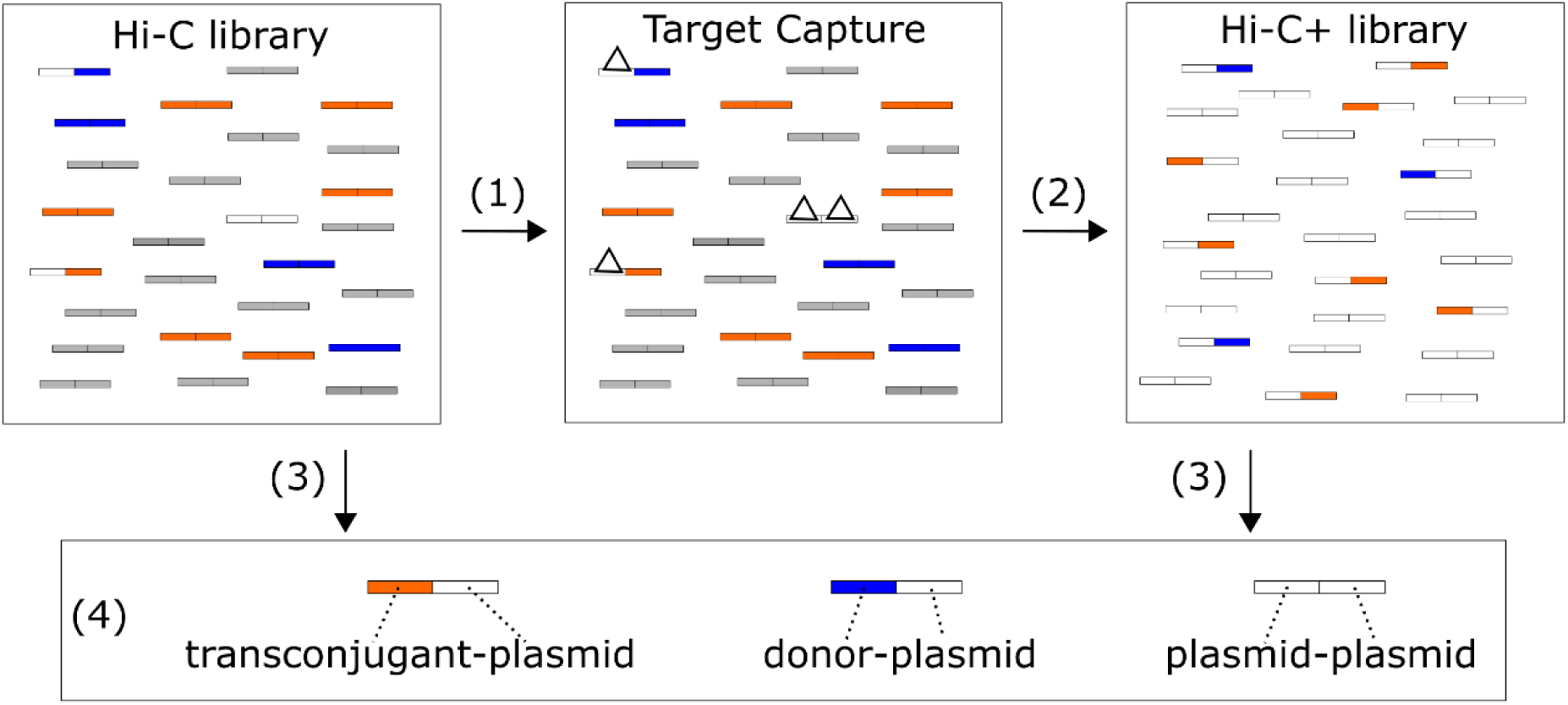
Overview of Hi-C+. 1) DNA from a Hi-C library was used as input for the target capture reaction. RNA baits complementary to the sequence of our plasmid (white triangles) were designed and applied to the Hi-C library. 2) The bait-bound library was purified and PCR-amplified, generating Hi-C+ libraries enriched for plasmid-specific DNA. 3) Hi-C and Hi-C+ libraries were sequenced. 4) Data analysis of these libraries consists of looking for all plasmid-associated Hi-C reads.

Comparison of the number of plasmid-associated reads (pB10-pB10, Ecoli-pB10 and Pputida-pB10) show that Hi-C+ enrichment was successful (Fig. 4A). Compared to Hi-C, the Hi-C+ libraries contained on average 3573 (SE ± 780) times more pB10-pB10 reads, 572 (SE ± 138) times more Ecoli-pB10 reads, and 343 (SE ± 186) times more Pputida-pB10 reads. The higher enrichment of pB10-pB10 reads was expected, as intracellular contacts between plasmid DNA fragments should be more frequent than contacts with the host chromosome. These fragments can also be bound by the RNA baits over their entire length, whereas roughly only half of Pputida-pB10 and Ecoli-pB10 Hi-C fragments can be bound by these baits. While the average enrichment was lower for Ecoli-pB10 and Pputida-pB10 Hi-C reads, it was still more than two orders of magnitude. This demonstrates that Hi-C+ can increase the proportion of plasmid DNA in Hi-C libraries.

**Fig. 4:**
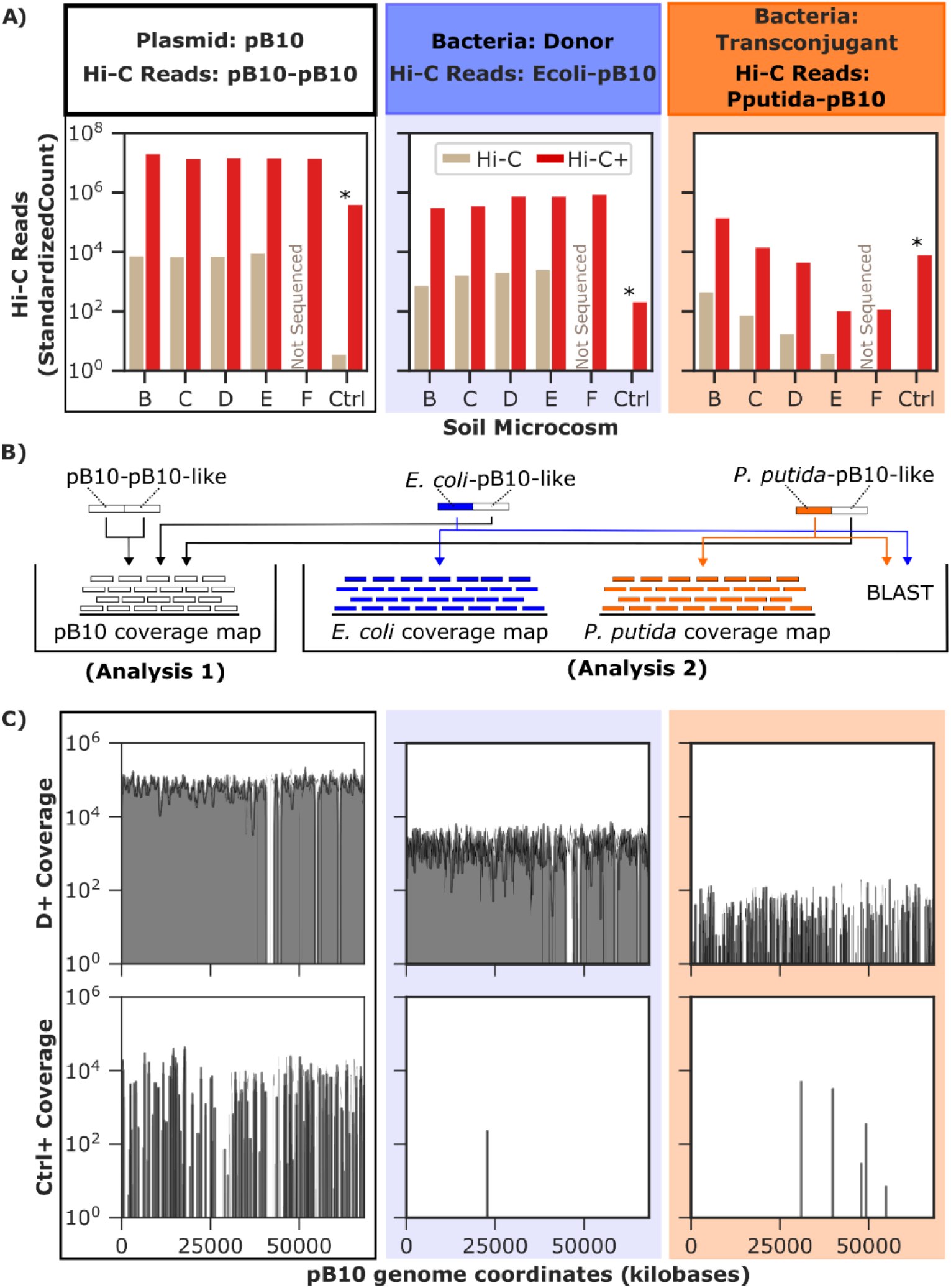
Detection of plasmid-linked reads in Hi-C+ libraries. A) Detection of pB10-pB10, Ecoli-pB10, and Pputida-B10 reads in each Hi-C and Hi-C+ library. *While the control microcosm had a high number of plasmid-associated eads, these were markedly different from those detected in other microcosms. B) To investigate this further, we carried ut an analysis on the plasmid-linked reads in the Control soil microcosm. Arrows demonstrate the analyses carried out n each of the Hi-C+ reads. Analysis 1 investigates where the pB10 segment of plasmid-like reads align on the plasmid eference genome. Analysis 2 determines where the *P. putida* KT2442 and *E. coli* MG1655 segments of plasmid-like eads align on their respective reference genomes. This is accompanied by a BLAST search of the sequence against the CBI nucleotide database to determine whether the region to which the reads align is conserved across bacteria. C) omparison of results from analysis 1 carried out on reads from a microcosm to which we added pB10 (D+) and the ontrol (Ctrl+). See Fig. S2 for all coverage maps. Gaps in coverage maps for D+ pB10-pB10 and Ecoli-pB10 Hi-C+ eads are areas of the pB10 genome that were identical to regions of the *P. putida* KT2442 or *E. coli* MG1655 genomes. esults from analysis (2) are shown in Fig. S3. Each column is color-coded to indicate the Hi-C reads for which data is resented.

### Target capture enriched genes commonly found in a soil microbiome

While Hi-C+ enriched pB10 plasmid DNA in the Hi-C libraries from soil microcosms B-F, we also observed an enrichment in the control soil microcosm that was not inoculated with pB10 (Fig. 4). This result suggests that members of the soil community contain DNA fragments identical to that of our plasmid and donor and recipient strains. Indeed, IncP1 plasmids and some pB10 genes such as transposons, integrons and resistance genes are ubiquitous in soil (*17*, *50*, *51*). Therefore, we hypothesized that in absence of pB10, target capture enriched sequences that are non-specific to our focal plasmid but commonly found in the soil bacterial community. We refer to these sequences as pB10-like in Fig. 4, to differentiate them from those observed in microcosms to which we added our focal plasmid.

Because the positive Hi-C+ signal in the negative control soil microcosm could interfere with the tracking of a focal plasmid, we explored these results in more detail. First, we looked at where the pB10-like reads aligned on the pB10 reference genome. This was done for all three types of plasmid-associated reads (Fig. 4B). Reads mapping to pB10 aligned to common genes amongst plasmids. For example, genes from the *trb* and *tra* operons (see Fig. S2 for all genes detected). Additionally, coverage provided by pB10-like reads in the control microcosm was sparser than a microcosm to which we added pB10 (Fig. 4C). The difference in coverage and detection of genes commonly found in soil indicate that those pB10-like reads are likely a reflection of the existing reservoir of mobile genetic elements in soil. Next, we examined the chromosomal side of Ecoli-pB10-like and Pputida-pB10-like Hi-C reads (Fig. 4B). Doing so can provide insight on whether these sequences are common amongst many bacteria. A BLAST search of the few *E. coli* and *P. putida* segments revealed that these were present across many bacteria within their respective genus (Fig. S3). This further reinforces the notion that the detection of plasmid-associated reads in our negative control is likely due to certain bacteria in soil that contain chromosomal genes like some in our donor and recipient, and plasmids or integrative conjugative elements with gene homologs like those encoded by pB10. Thus, despite some background enrichment by Hi-C+, it is possible to distinguish noise from focal plasmid-host links by using our coverage metrics.

### Exploitation of plasmid coverage and uniqueness helps discriminate specific Hi-C+ reads from background noise and non-specific target

The detection of pB10-like reads in our control soil microcosm indicates that specificity of the target capture reaction is affected by the presence of targets similar to our focal plasmid, and additional steps are needed to discriminate pB10-specific from non-specific enriched sequences. We therefore incorporate two key metrics: plasmid coverage and unique plasmid bases. First, if a target capture reaction specifically enriches a Hi-C library with our focal plasmid pB10, then we expect that those Hi-C+ reads map over the entire length of the plasmid sequence. This would result in a higher and more even plasmid coverage than a non-specific enrichment reaction, which would be more scattered or specific to a few locations on the plasmid map (see Fig. 4C). Second, the inclusion of a control microcosm enabled us to identify the segments unique to pB10. At each location in the pB10 sequence, we compared whether that base was detected in sequencing data for the control microcosm. Locations that were not detected in the control microcosm were denoted ‘unique plasmid bases’ and were likely specific to our focal plasmid. We then calculated the percentage of these unique plasmid bases. Similarly to plasmid coverage, we expect that successful enrichment result in a higher percentage of unique plasmid bases detected in the treated soils. In contrast, a reaction that enriches for pB10-like DNA will contain the same sequences as the control microcosm and will therefore have a low unique plasmid base percentage. Using this percentage in combination with the plasmid coverage can facilitate differentiation between specific and nonspecific target-capture reactions.

We applied our plasmid coverage metrics to the paired-end Hi-C reads from each soil microcosm. Consistent with our expectations, pB10-pB10 Hi-C reads and those indicating the presence of the donors (Ecoli-pB10) had a high plasmid coverage and percentage of unique plasmid bases detected in every pB10-treated soil microcosm (Fig. 5). This was the case for Hi-C alone and Hi-C+. For reads indicating the presence of transconjugants (Pputida-pB10), the Hi-C+ libraries from microcosms B, C, D, and F showed high plasmid coverage and percentage of unique plasmid bases (Fig. 5). Most importantly, in the control, the coverage was much lower than the microcosms to which we added pB10 (Fig. 5). This highlights that focal plasmid coverage, especially intra-plasmid Hi-C reads, can be used to assess whether a plasmid of interest was successfully enriched for.

**Fig. 5:**
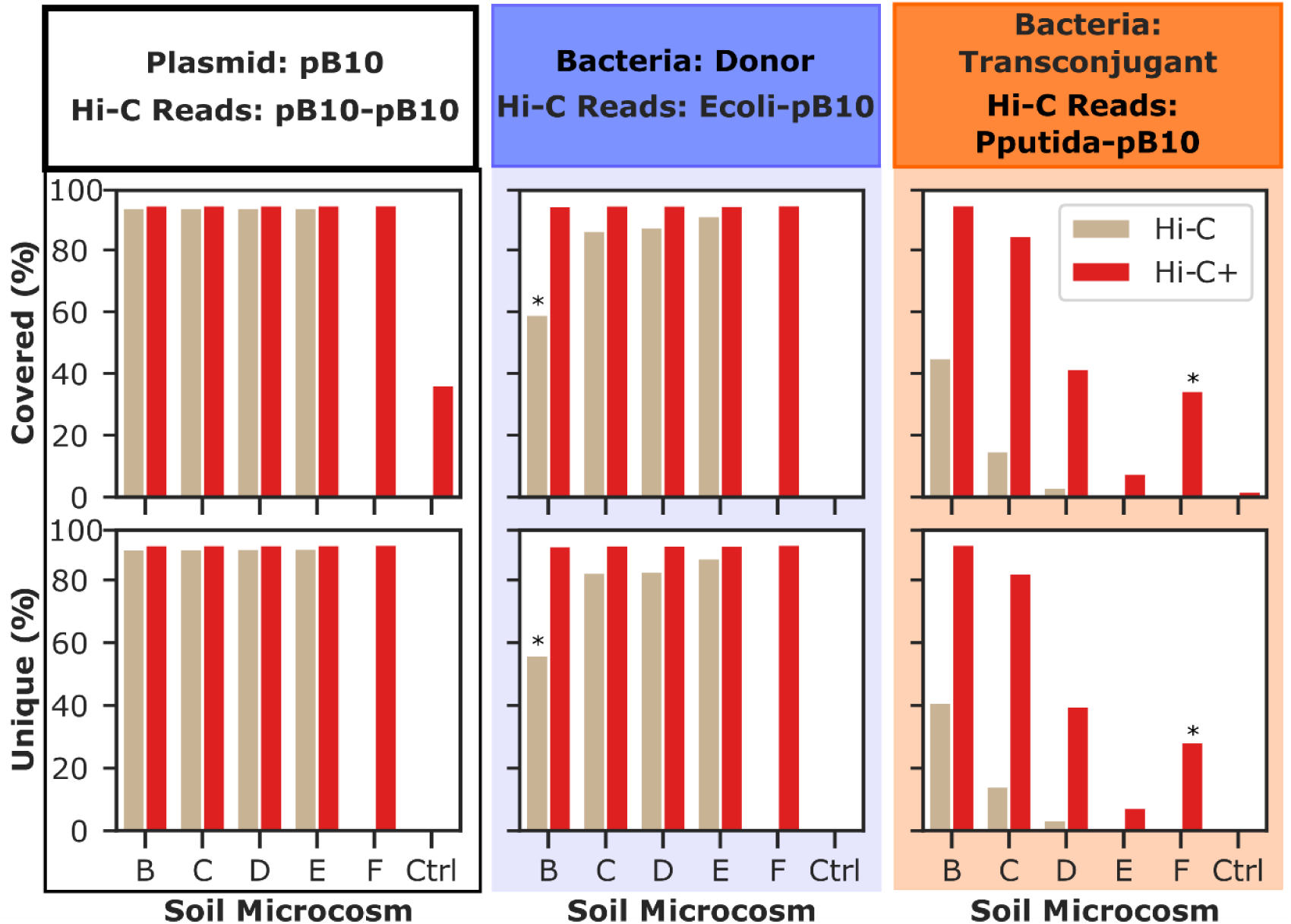
Plasmid coverage and unique bases help identify specific target capture reactions. Specific enrichment of a focal plasmid in the treated microcosms can be discerned by comparing the percentage of plasmid coverage (Top) and unique plasmid bases captured (Bottom) to the control microcosm. Each column is color-coded to indicate the Hi-C and Hi-C+ reads for which data is presented. *Two deviations from our expectations were observed in our results. The first is the lower score for Hi-C donor reads in microcosm B. We expect this is a result of there being a higher amount of transconjugant than donors in this microcosm, which affected their proportions in the sequenced library. The second is the higher scores for transconjugant Hi-C+ reads in microcosm F compared to E. We expect this occurred due to differences in the extraction efficiency of the bacterial portion from soil (see methods) coupled with experimental variation in target capture.

### Hi-C+ can identify the new hosts of a focal plasmid in soil

In our experimental design, we knew the identity of the transconjugant, *P. putida* KT244 (pB10) and had its reference sequence, which facilitated its detection in each of our soil microcosms. However, researchers seeking to use Hi-C+ to monitor the transfer of a plasmid from a donor strain to the members of any microbiome will not have a reference sequence for their unknown new plasmid hosts. Instead, they will have to use taxonomic classifiers to identify the putative new hosts of their focal plasmid. We sought to emulate this process by attempting a blind *de novo* identification of our *P. putida* KT2442 transconjugant. An overview of the data processing pipeline is shown in Fig. 6A. Only reads that consisted of pB10 sequence on one side and unknown sequence on the other, designated “pB10-unaligned”, were retained. For the assignment of taxonomic classification, the unaligned portion of these reads were used as input for Kraken2 (*52*). This tool uses k-mers within each sequence to map the unaligned read to the lowest common ancestor within the standard Kraken2 database. For each taxonomic rank, classified reads were used to calculate the plasmid coverage metrics established in the previous section (plasmid coverage and percentage of unique plasmid bases detected) (Fig. 6A).

**Fig. 6:**
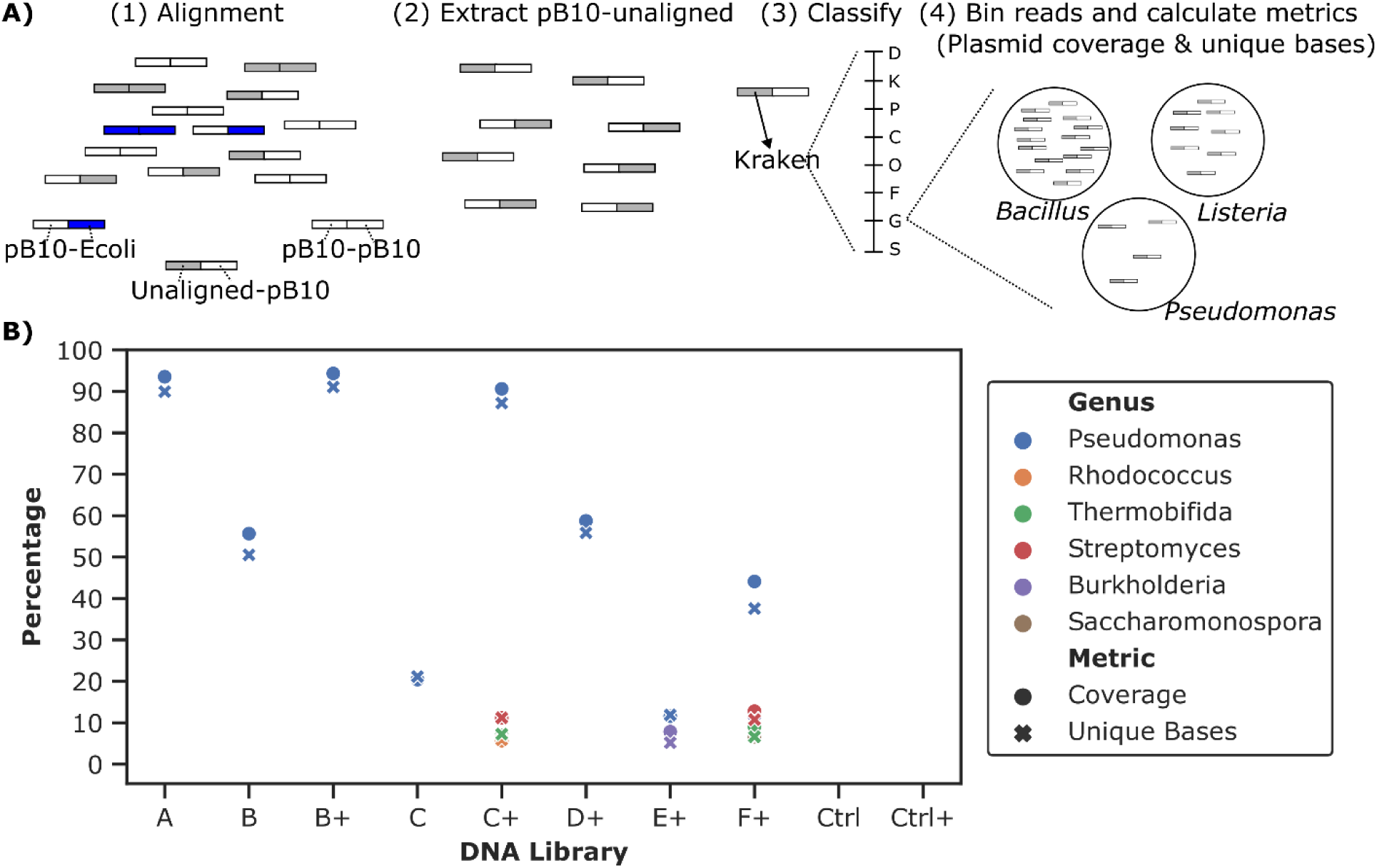
*De novo* identification of pB10 hosts. A) Overview of data processing pipeline. 1) Reads are first aligned only to pB10::rfp and *E. coli* MG1655 reference sequences, generating the plasmid-associated Hi-C reads depicted at the bottom. 2) pB10-unaligned reads are extracted. 3) The unaligned portion of pB10-unaligned reads is used to assign taxonomic classification using Kraken2. Classification is assigned at each taxonomic rank, from domain (D) to species (S). 4) Read binning and calculation of genome coverage metrics (plasmid coverage and percent unique plasmid bases). An example of read binning at the genus level is shown. Reads are binned within each classification, here *Bacillus*, *Listeria* or *Pseudomonas,* and the genome coverage metrics are calculated for each bin. B) Genus level identification of pB10 hosts and associated metrics. The plus signs in the X axis differentiate results from standalone Hi-C libraries, and the Hi-C+ libraries to which we applied target capture. Genera with less than 5% plasmid coverage and unique plasmids bases were excluded from the visualization. Except for the controls, the libraries with no genus above the established cutoff were also excluded from visualization.

We applied this workflow to the Hi-C and Hi-C+ reads from each of our soil microcosms. In libraries A, B+ and C+, we were able to identify *Pseudomonas* as the transconjugant with high confidence. That is, reads linking pB10 to this genus had high scores for the two metrics we proposed, plasmid coverage and unique plasmid bases detected (Fig. 6B). The *Pseudomonas* genus was also identified as the transconjugant in libraries B, C, D+, and F+, although with lower percentage of plasmid coverage and unique plasmid bases detected. Notably, below 20%, we identified numerous other genera as transconjugants. These hosts are likely false positives that result from bacteria in soil that carry plasmids similar to our focal plasmid and thus achieve low scores for our metrics. We propose that transconjugants with scores below 20% are disregarded as noise and that scores above this threshold be interpreted with increasing confidence as the genome coverage metrics increase. Unsurprisingly, we were unable to confidently identify the host of the plasmid down to the species level, *P. putida*. Identification with high confidence below the genus level seems unlikely due the short sequence (less than 200 bp) and the high similarity between the genomes of species within a genus.

## Discussion

Plasmids are important mediators of horizontal gene transfer in microbial communities (*53*). Across multiple habitats they play a crucial role in mobilizing antibiotic resistance genes, which can eventually transfer to human pathogens (*16*). It is therefore pivotal to detect plasmid transfer and identify new plasmid hosts in natural communities and complex matrices. To this end, we developed Hi-C+, a method that can identify rare plasmid hosts. In this study, we used Hi-C+ to identify transconjugants resulting from the transfer of a focal plasmid in a microbial community. However, its application is not limited to this scenario. It can also be employed for identifying the host range of natural plasmids, mobile genetic elements, or genes encoding a particular phenotype under study. The implementation of Hi-C+ can provide important insights into the key players and factors that drive the vertical and horizontal transfer of antibiotic resistance genes, which can in turn be used to design strategies to limit their spread.

We show that the standalone Hi-C method can be used to link a focal plasmid to its hosts in a diverse microbiome such as an agricultural soil. While Hi-C has previously been used to identify plasmid hosts (*40–43*, *54*), a quantitative assessment of its detection limit hasn’t been carried out. Here, we determine that plasmid hosts can be identified even when present at densities as low as 10^-5^ or 0.001% of the total bacterial community. Moreover, our findings suggest that Hi-C can discriminate rare transconjugants in the presence of the donors and the natural soil community and can even help identify the host taxon when combined with a target enrichment approach. Additionally, we defined a model that can be used to relate Hi-C reads to bacterial counts in cfu/g. While the quantitative aspect of this method is limited to plasmid hosts present above a relative abundance of 10^-5^ and Hi-C datasets prior to application of target capture, it provides a way to obtain quantitative information on plasmid containing cells. Such information can be useful for understanding the ecology of plasmids in natural habitats.

Our Hi-C+ approach increased the detection limit of the Hi-C method for associating a plasmid with its hosts in soil by using target enrichment. The degree of enrichment varied between targets, with the plasmid donors being enriched on average 500-fold and the transconjugants 300-fold. These findings are comparable with those from previous studies employing target capture (*55*, *55*, *56*). For example, one study found an increase in read counts of more than two orders of magnitude for 31 of their 32 targets (*57*). We observed a difference in degree of transconjugant enrichment across microcosms, with it decreasing as the number of added transconjugants decreased. We expect that this difference is due to the higher abundance of donors in most of our soil microcosms. In our target capture reactions, donor and transconjugant Hi-C reads are in competition for plasmid baits. The higher abundance of donor reads in most of the soil microcosm Hi-C libraries likely resulted in their more successful enrichment, which is further exacerbated by the use of multiple PCR cycles in the protocol. This highlights that, when using Hi-C+ for studying transfer of a focal plasmid, the original plasmid hosts (donors) need to drop in abundance below that of transconjugants to limit the extent to which they sequester the plasmid baits in the reaction. In sum, the combination of target capture with Hi-C enabled us to improve the detection limit of the method for linking a plasmid to its host.

We also showed that Hi-C+ can be used to identify the host of a focal plasmid in soil at the genus level. When attempting to identify the transconjugant in our set-up without using any alignment method, we were able to do so at the genus level (Pseudomonas) when the relative abundance of this strain in the soil community was as low as at 10^-7^. This limit corresponds to soil microcosm F, where reads linking pB10 to the *Pseudomonas* genus had >30% plasmid coverage and unique plasmid bases detected. Importantly, this represents a significant improvement compared to Hi-C alone. Similar confidence in host identification with Hi-C was only achieved in soil microcosm A, B and C, where the transconjugants were respectively present at 10^-1^, 10^-2^, and 10^-3^ of the total bacterial community. This demonstrates that, along with increasing the detection limit of Hi-C, Hi-C+ also significantly improves our ability to identify new transconjugants.

Lastly, we identified several key limitations that need to be considered in future microbiome studies that use target capture on Hi-C reads. As observed in our control soil microcosm, enrichment for similar fragments of DNA in the microbiome under study can cause non-specific enrichment (here pB10-like reads), hindering interpretation of Hi-C+ data. To circumvent this limitation, a possible approach is the exclusion of regions commonly shared between plasmids in the probe design, e.g., accessory genes such as those encoding resistance, transposases, or conjugative transfer. One could also leverage existing knowledge of the microbiome under study to exclude specific areas of the focal plasmid. In our application, this could be the exclusion of regions common to IncP plasmids that are typically present in soil (*51*, *58*, *59*). In cases where researchers would rather not remove probes, we have proposed the inclusion of two plasmid coverage metrics to increase confidence in putative hosts identified by Hi-C+. The first is to determine whether the reads linking a focal plasmid to a host cover the entire length of the focal plasmid sequence. The second is calculating the percentage of unique plasmid bases detected. This requires that a target capture reaction be carried out on a control microbiome sample that does not contain the focal plasmid. Subsequently, we can identify which areas of the focal plasmid are identical to that of other members of the microbiome and rule out any probe binding to those regions. Using these metrics increases confidence regarding the identification of plasmid hosts.

Despite the fact that many microbes occur at low frequencies within their community, the rare biosphere remains an unexplored and often overlooked part of microbiomes (*60*). These rare microbes may have critical roles in the ecology of microbial communities. For example, they function as a refugia where genetic information is archived and can be accessed and selected when environmental conditions change (*61*), such as in response to the presence of antibiotics or heavy metals. Here, we demonstrate the application of Hi-C and Hi-C+ in soil microcosms to detect and identify the host of a focal plasmid. We show that this approach can identify the host of a plasmid even when present in low abundance in a community, and thereby provide new insights into rare plasmid reservoirs in diverse habitats.

## Materials and Method

### Strains and soil

Soil was obtained from the USDA ARS Northwest Irrigation and Soils Research Laboratory in Kimberly, ID and stored at 4 °C until the time of the experiment. The three strains used in our study were the plasmid donors *Escherichia coli* K-12 MG1655Nal::gfp (pB10) (*62*), the plasmid-free recipients *Pseudomonas putida* KT2442, and the transconjugants *P. putida* KT2442 (pB10) (*63*, *64*). The strains and reference genomes were obtained from previous studies (*62–64*). The reference for *E. coli* K12 MG1655 was manually modified to include the mini-*Tn*5-*gfp* transposon. Reads from a sequenced *E. coli* K12 MG1655 were mapped to the transposon reference. Subsequently, the overhanging reads were used to identify the location on the *E. coli* MG1655 sequence that the transposon had inserted and the reference was manually edited. The same process was carried out for annotating the pB10 sequence with the mini-*Tn5*-rfp transposon.

All three strains were grown overnight in lysogeny broth (LB; Thermo Fisher Scientific, Waltham, MA, USA) with shaking at 200 r.p.m. The donor was grown at 37 °C in broth containing nalidixic acid (50 mg. L^-1^) and kanamycin (50 mg. L^-1^). The recipient was grown at 30 °C in broth containing rifampicin (100 mg. L^-1^), and the transconjugant in broth containing rifampicin (100 mg. L^-1^) and kanamycin (50 mg. L^-1^).

### Set up of soil microcosms

Our soil microcosms were set up to simulate plasmid transfer scenarios in natural soil. This was accomplished by adding mixtures of donors, recipients, and transconjugants to 6 grams of soil in 50-mL Falcon tubes (see Table 1). To determine the number of bacteria to be added to each soil microcosm, we first estimated the total number of soil bacteria by quantifying the copy number of the 16S rRNA gene by qPCR. This protocol was followed as described by Hill *et al* (*65*), using the primers designed by Liu *et al* (*66*). We used the 16S rRNA copies to estimate the bacteria per g of soil, assuming an average of 4.2 copies per bacterial chromosome (*67*). Using this value, 5.28 x 10^8^ bacteria/g of soil, we estimated the number of donors, recipients, and transconjugants to be added to each soil microcosm to obtain the relative abundances shown in Table S1. We then estimated the densities of donor, recipient, and transconjugant in the overnight cultures through plating on selective media and used these to determine the volume of cell suspensions to be added to each soil microcosm.

After overnight growth, the cultures were centrifuged for 3 min at 5000 x *g* and pellets were resuspended in 1/10^th^ volume of phosphate-buffered saline pH 7.4 (PBS) (Thermo Fisher Scientific, Waltham, MA, USA). For each soil microcosm (Table S1) the appropriate volume of each cell suspension was added into a tube and supplemented with PBS to a total volume of 150 µL. In a 50 mL falcon tube, 6 *g* of soil was mixed with this 150 µL mixture of donors, recipients, and transconjugants. The soil was thoroughly mixed with a sterile spatula and we immediately proceeded with preparation of Hi-C libraries.

### Sample processing and Hi-C library preparation

Prior to undergoing the Hi-C library preparation, the bacterial fraction was extracted and purified from the soil for each microcosm as follows. The soil was resuspended in 27 mL of sterile 2% sodium hexametaphosphate (Sigma-Aldrich, St. Louis, MO, USA). Samples were vortexed for 5 minutes, then serially diluted and plated on selective media to obtain the plate count data shown in Table 1 and Fig. 1. Subsequently, the vortexed samples underwent centrifugation at 150 x *g* for 30 sec. The supernatant was recovered into a new 50 mL falcon tube and centrifuged at 10,000 x *g* for 10 minutes. The resulting pellet was resuspended in 6 mL of Tris-Buffered Saline (TBS) (Thermo Fisher Scientific, Waltham, MA, USA), placed on a 5 mL OptiPrep^TM^ cushion (Sigma-Aldrich, St. Louis, MO, USA), and centrifuged for 20 minutes at 10,000 x *g*. The top layer from the density gradient was recovered and centrifuged for 10 minutes at 17,000 x *g*. The pellet was then resuspended in TBS and aliquoted into five micro-centrifuge tubes. Each aliquot was centrifuged for 5 minutes at 17,000 x *g*, the pellet was then resuspended in 1 mL of cross-linking solution (1% formaldehyde (Thermo Fisher Scientific, Waltham, MA, USA)) and incubated for 20 minutes at room temperature. Then, 100 µL of quenching solution (125 mM glycine (Sigma-Aldrich, St. Louis, MO, USA) was added to each sample, followed by incubation for 20 more minutes. Samples were centrifuged at 17,000 x *g* for 5 minutes and the supernatant was discarded. The resulting pellet was washed with 1 mL of TBS, centrifuged at 17,000 x *g* for 5 minutes, and stored at −20 °C until library preparation. To avoid experimental variation, all soil microcosms were set up and processed at the same time. The cross-linked samples were sent to Phase Genomics^©^ for preparation of Hi-C libraries using their ProxiMeta^TM^ kit according to the associated protocol.

### Preparation of Hi-C+ libraries

In this study we developed the combination of Hi-C with target capture and named it Hi-C+. This method consists of the enrichment of target DNA within the Hi-C libraries using the myBaits^®^ (Daicel Arbor Biosciences, Ann Arbor, MI, USA) custom target enrichment protocol. Design of probes was carried out by Daicel Arbor Biosciences, with 70mer probe length and 20 bp tiling over the entire length of the genome sequence of plasmid pB10. Probes were screened against the *E. coli* K-12 MG1655 and *P. putida* KT2442 reference sequences. Those with high similarity to the references were removed, along with probes that had a significant percent of simple repeats or low-complexity DNA (>50% soft-repeat-masking, screened by Daicel Arbor Biosciences using RepeatMasker). This resulted in 3220 unique probes.

Hi-C libraries were first pre-treated for compatibility with myBaits^®^ target enrichment kit. The ProxiMeta^TM^ kits that are employed by Phase Genomics^©^ use Nextera^TM^ technology to prepare libraries for sequencing. These are incompatible with myBaits^®^ target enrichment kit and the remaining streptavidin-affinity molecules must be depleted. The depletion reaction was carried out using 200 ng of library DNA as input. Each library underwent four rounds of PCR amplification with KAPA HiFi HotStart ReadyMix (Roche Holding AG, Basel, CH) and universal primers for amplifying Nextera libraries. The amplification reactions were pelleted, resuspended in 30 µL of Dynabeads^TM^ MyOne^TM^ Streptavidin C1 beads (Invitrogen, Waltham, MA, USA), and pelleted on a magnetic particle concentrator. The supernatant was removed and cleaned using Zymo Research^©^ DNA Clean and Concentrator kit (Zymo Research^©^, Irivine, CA, USA). The resulting cleaned DNA was used as input for the target enrichment reaction.

The target capture reactions were carried out using the myBaits “v4” chemistry and associated v4.01 manual. Since the plasmid DNA makes up a small fraction of our libraries, we used a modified high-sensitivity protocol. The temperature for hybridization and washing steps was changed to 62 °C. In the first enrichment (step 1.2.2 in manual), we used 4.4 µL of baits and 1.1 µL of nuclease-free water. After the first enrichment, two amplifications per capture reaction were carried out (step 3.3 in the manual). The purified PCR products were combined, concentrated to 7 µL using Zymo Research^©^ DNA Clean and Concentrator kit, and used as input for a second round of target capture. For the second enrichment (step 1.2.2 in manual), we used 1.1 µL of baits and 4.4 µL of nuclease-free water. All other steps were carried out as instructed in the v4.01 manual.

### Sequencing of Hi-C and Hi-C+ libraries

Hi-C and Hi-C+ library size selection, quantification, and pooling were carried out by the IIDS Genomics and Bioinformatics Resources Core at the University of Idaho (Moscow, ID, USA). The libraries were then sequenced at the University of Oregon sequencing core (Eugene, OR, USA). The six Hi-C libraries were sequenced using two lanes of HiSeq 4000, 2 x 100 bp paired-end, while the six Hi-C+ libraries were sequenced on three lanes of NovaSeq 6000, 2 x 150 bp paired-end reads. The number of paired-end reads obtained from each library was as follows; A: 102,818,926, B: 97,688,195, C: 155,702,732, D: 127,023,749, E: 146,798,824, Ctrl1: 103,992,690, B+: 67,166,513, C+: 466,371,775, D+: 56,882,152, E+: 83,617,103, F+: 405,252,012, Ctrl1+: 58,053,294, Ctrl2+: 38,173,655.

### Data Analysis

The sequencing reads first underwent read trimming and filtering using fastp with default arguments and minimum length requirement of 50 base pairs (*68*). We then aligned our data to our reference sequences using BWA Mem with the -5SPY options (*69*). Hi-C and Hi-C+ read counts were subsequently analyzed using a custom python script. In this script, we binned together all the paired-end reads for which at least one end mapped to the sequence of pB10 and the other end mapped the sequence of *E. coli* MG1655, *P. putida* KT2442, or pB10. Two important criteria were defined for including alignments in these bin. First, only reads that aligned to the reference genome over their entire length and had 100% nucleotide identity were used in the analysis. Second, we only retained alignments that were mapped to a single reference. Multi-mapped reads that aligned equally well to multiple reference genomes were excluded to prevent false enrichment signals from similar DNA on the plasmid and bacterium sequences. While the probe design was intended to limit this occurrence, it was carried out on reference genomes that did not contain the mini-transposons with *gfp* in our marked *E. coli* MG1655 Nal::gfp and *rfp* in pB10. As a result, there were probes that bind to both the pB10 and *E. coli* genomes, specifically the mini-transposon used to insert the fluorescent protein genes. All scripts and detailed descriptions are available on GitHub (https://github.com/scastanedabarba/hic_targetcapture).

### Mathematical Models

The model for predicting bacterial count was fitted using the glm.nb function in the MASS v7.3-58.2 R package. Our model is a negative binomial regression (Equation 1) with a response variable of bacterial count (cfu/g) and predictor variables Pputida-pB10 Hi-C reads and sequencing depth.

### De Novo taxonomic classification of pB10 hosts

Trimmed reads were first aligned only to the *E. coli* MG1655 and pB10 sequences using BWA Mem with the -5SPY options (*69*). After alignment, a custom python script was used to extract paired-end reads whereby one pair was aligned to pB10 and the other was unaligned. The same criteria defined for filtering alignments in our ‘data analysis’ section were used here. These represent cross-linked DNA between the plasmid and putative hosts. The unaligned portion of these reads were output to new FASTA files and underwent taxonomic classification using Kraken2 with minimum confidence score of .1, using the standard database (*52*). Taxonkit and csvtk were then used to convert taxids to full taxonomic lineages (*70*), followed by use of a python script to calculate metrics for each classification, at each taxonomic rank.

## Acknowledgements

Data collection and analyses performed by the Institute for Interdisciplinary Data Sciences (IIDS) Genomics Resources Core at the University of Idaho were supported in part by NIH COBRE grant P30GM103324. The authors would like to thank Robert Dungan from the USDA ARS Northwest Irrigation and Soils Research Laboratory in Kimberly, ID for help in acquiring soil. The authors would also like to thank Olivia Kosterlitz, Erin Mack, and Morgan Sower for critical reading of the manuscript.

## Funding

National Institute of Food and Agriculture (NIFA) grant no. 2018-67017-27630 of the United States Department of Agriculture (E.M.T., T.S., B.J.R.)

Bioinformatics and Computational Biology Program at the University of Idaho in partnership with the Institute for Bioinformatics and Evolutionary Studies (now Institute for Interdisciplinary Data Sciences, IIDS). (S.C.B)

## Author Contributions

E.M.T. and T.S. conceived the project; E.M.T, T.S. and S.C. designed the study; S.C. performed the experiments, collected the data, and performed data analysis; S.C. and B.J.R. designed the math model; T.S., E.M.T., and S.C. wrote the paper. All authors helped revise the paper.

## Competing Interests

The authors declare no competing interests.

## Data and materials availability

All sequencing data pertaining to this project have been made available at the National Center for Biotechnology Information Sequencing Read Archive (PRJNA1040204).

## Notes

### Competing Interest Statement

The authors have declared no competing interest.

## References

1. S. Redondo-Salvo, R. Fernández-López, R. Ruiz, L. Vielva, M. de Toro, E. P. C. Rocha, M. P. Garcillán-Barcia, F. de la Cruz, Pathways for horizontal gene transfer in bacteria revealed by a global map of their plasmids. Nat. Commun. 11, 3602 (2020).

2. L. S. Frost, R. Leplae, A. O. Summers, A. Toussaint, Mobile genetic elements: the agents of open source evolution. Nat. Rev. Microbiol. 3, 722–732 (2005).

3. J. P. J. Hall, M. A. Brockhurst, E. Harrison, Sampling the mobile gene pool: innovation via horizontal gene transfer in bacteria. Philos. Trans. R. Soc. 372, 20160424 (2017).

4. A. Norman, L. H. Hansen, S. J. Sørensen, Conjugative plasmids: vessels of the communal gene pool. Philos. Trans. R. Soc. 364, 2275–2289 (2009).

5. C. J. Murray, K. S. Ikuta, F. Sharara, L. Swetschinski, G. R. Aguilar, A. Gray, C. Han, C. Bisignano, P. Rao, E. Wool, S. C. Johnson, A. J. Browne, M. G. Chipeta, F. Fell, S. Hackett, G. Haines-Woodhouse, B. H. K. Hamadani, E. A. P. Kumaran, B. McManigal, R. Agarwal, S. Akech, S. Albertson, J. Amuasi, J. Andrews, A. Aravkin, E. Ashley, F. Bailey, S. Baker, B. Basnyat, A. Bekker, R. Bender, A. Bethou, J. Bielicki, S. Boonkasidecha, J. Bukosia, C. Carvalheiro, C. Castañeda-Orjuela, V. Chansamouth, S. Chaurasia, S. Chiurchiù, F. Chowdhury, A. J. Cook, B. Cooper, T. R. Cressey, E. Criollo-Mora, M. Cunningham, S. Darboe, N. P. J. Day, M. D. Luca, K. Dokova, A. Dramowski, S. J. Dunachie, T. Eckmanns, D. Eibach, A. Emami, N. Feasey, N. Fisher-Pearson, K. Forrest, D. Garrett, P. Gastmeier, A. Z. Giref, R. C. Greer, V. Gupta, S. Haller, A. Haselbeck, S. I. Hay, M. Holm, S. Hopkins, K. C. Iregbu, J. Jacobs, D. Jarovsky, F. Javanmardi, M. Khorana, N. Kissoon, E. Kobeissi, T. Kostyanev, F. Krapp, R. Krumkamp, A. Kumar, H. H. Kyu, C. Lim, D. Limmathurotsakul, M. J. Loftus, M. Lunn, J. Ma, N. Mturi, T. Munera-Huertas, P. Musicha, M. M. Mussi-Pinhata, T. Nakamura, R. Nanavati, S. Nangia, P. Newton, C. Ngoun, A. Novotney, D. Nwakanma, C. W. Obiero, A. Olivas-Martinez, P. Olliaro, E. Ooko, E. Ortiz-Brizuela, A. Y. Peleg, C. Perrone, N. Plakkal, A. Ponce-de-Leon, M. Raad, T. Ramdin, A. Riddell, T. Roberts, J. V. Robotham, A. Roca, K. E. Rudd, N. Russell, J. Schnall, J. A. G. Scott, M. Shivamallappa, J. Sifuentes-Osornio, N. Steenkeste, A. J. Stewardson, T. Stoeva, N. Tasak, A. Thaiprakong, G. Thwaites, C. Turner, P. Turner, H. R. van Doorn, S. Velaphi, A. Vongpradith, H. Vu, T. Walsh, S. Waner, T. Wangrangsimakul, T. Wozniak, P. Zheng, B. Sartorius, A. D. Lopez, A. Stergachis, C. Moore, C. Dolecek, M. Naghavi, Global burden of bacterial antimicrobial resistance in 2019: a systematic analysis. Lancet 399, 629–655 (2022).

6. F. Sun, D. Zhou, Q. Sun, W. Luo, Y. Tong, D. Zhang, Q. Wang, W. Feng, W. Chen, Y. Fan, P. Xia, Genetic characterization of two fully sequenced multi-drug resistant plasmids pP10164-2 and pP10164-3 from *Leclercia adecarboxylata*. Sci. Rep. 6, 33982 (2016).

7. L. Falgenhauer, H. Ghosh, S. Doijad, Y. Yao, B. Bunk, C. Spröer, M. Kaase, R. Hilker, J. Overmann, C. Imirzalioglu, T. Chakraborty, Genome analysis of the carbapenem-and colistin-resistant *Escherichia coli* isolate NRZ14408 reveals horizontal gene transfer pathways towards panresistance and enhanced virulence. Antimicrob. Agents Chemother. 61 (2017).

8. L. Falgenhauer, S.-E. Waezsada, Y. Yao, C. Imirzalioglu, A. Käsbohrer, U. Roesler, G. B. Michael, S. Schwarz, G. Werner, L. Kreienbrock, T. Chakraborty, RESET consortium, Colistin resistance gene *mcr-1* in extended-spectrum *β-lactamase*-producing and carbapenemase-producing Gram-negative bacteria in Germany. Lancet. Infect. Dis. 16, 282– 283 (2016).

9. J. R. Mediavilla, A. Patrawalla, L. Chen, K. D. Chavda, B. Mathema, C. Vinnard, L. L. Dever, B. N. Kreiswirth, Colistin-and carbapenem-resistant *Escherichia coli* harboring *mcr-1* and *blaNDM-5*, causing a complicated urinary tract infection in a patient from the United States. mBio 7, e01191–16 (2016).

10. P. McGann, E. Snesrud, R. Maybank, B. Corey, A. C. Ong, R. Clifford, M. Hinkle, T. Whitman, E. Lesho, K. E. Schaecher, *Escherichia coli* harboring *mcr-1* and *blaCTX-M* on a novel IncF plasmid: First report of *mcr-1* in the United States. Antimicrob. Agents Chemother. 60, 4420–4421 (2016).

11. B. McCollister, C. V. Kotter, D. N. Frank, T. Washburn, M. G. Jobling, Whole-genome sequencing identifies *in vivo* acquisition of a *blaCTX-M-27*-carrying IncFII transmissible plasmid as the cause of ceftriaxone treatment failure for an invasive *Salmonella enterica Serovar Typhimurium* Infection. Antimicrob Agents Chemother 60, 7224–7235 (2016).

12. M. Zhuang, Y. Achmon, Y. Cao, X. Liang, L. Chen, H. Wang, B. A. Siame, K. Y. Leung, Distribution of antibiotic resistance genes in the environment. Environ. Pollut. 285, 117402 (2021).

13. L. Poirel, V. Cattoir, P. Nordmann, Plasmid-mediated quinolone resistance; Interactions between human, animal, and environmental ecologies. Front. Microbiol. 3 (2012).

14. R. Wang, L. van Dorp, L. P. Shaw, P. Bradley, Q. Wang, X. Wang, L. Jin, Q. Zhang, Y. Liu, A. Rieux, T. Dorai-Schneiders, L. A. Weinert, Z. Iqbal, X. Didelot, H. Wang, F. Balloux, The global distribution and spread of the mobilized colistin resistance gene *mcr-1*. Nat. Commun. 9, 1179 (2018).

15. Y.-Y. Liu, Y. Wang, T. R. Walsh, L.-X. Yi, R. Zhang, J. Spencer, Y. Doi, G. Tian, B. Dong, X. Huang, L.-F. Yu, D. Gu, H. Ren, X. Chen, L. Lv, D. He, H. Zhou, Z. Liang, J.-H. Liu, J. Shen, Emergence of plasmid-mediated colistin resistance mechanism MCR-1 in animals and human beings in China: a microbiological and molecular biological study. Lancet Infect. Dis. 16, 161–168 (2016).

16. S. Castañeda-Barba, E. M. Top, T. Stalder, Plasmids, a molecular cornerstone of antimicrobial resistance in the One Health era. Nat. Rev. Microbiol., 1–15 (2023).

17. K. Blau, A. Bettermann, S. Jechalke, E. Fornefeld, Y. Vanrobaeys, T. Stalder, E. M. Top, K. Smalla, The transferable resistome of produce. mBio 9, e01300–18 (2018).

18. J. P. J. Hall, A. J. Wood, E. Harrison, M. A. Brockhurst, Source-sink plasmid transfer dynamics maintain gene mobility in soil bacterial communities. Proc. Natl. Acad. Sci. U.S.A. 113, 8260–8265 (2016).

19. W. Loftie-Eaton, A. Crabtree, D. Perry, J. Millstein, J. Baytosh, T. Stalder, B. D. Robison, L. J. Forney, E. M. Top, Contagious antibiotic resistance: Plasmid transfer among bacterial residents of the Zebrafish gut. Appl. Environ. Microbiol. 87, e02735–20 (2021).

20. R. Marti, A. Scott, Y.-C. Tien, R. Murray, L. Sabourin, Y. Zhang, E. Topp, Impact of manure fertilization on the abundance of antibiotic-resistant bacteria and frequency of detection of antibiotic resistance genes in soil and on vegetables at harvest. Appl. Environ. Microbiol. 79, 5701–5709 (2013).

21. L. Pérez-Etayo, D. González, J. Leiva, A. I. Vitas, Multidrug-Resistant Bacteria Isolated from Different Aquatic Environments in the North of Spain and South of France. Microorganisms 8, 1425 (2020).

22. D. Versluis, T. de J. B. González, E. G. Zoetendal, M. W. J. van Passel, H. Smidt, High throughput cultivation-based screening on porous aluminum oxide chips allows targeted isolation of antibiotic resistant human gut bacteria. PLOS ONE 14, e0210970 (2019).

23. V. H. T. Pham, J. Kim, Cultivation of unculturable soil bacteria. Trends Biotechnol 30, 475– 484 (2012).

24. U. Klümper, L. Riber, A. Dechesne, A. Sannazzarro, L. H. Hansen, S. J. Sørensen, B. F. Smets, Broad host range plasmids can invade an unexpectedly diverse fraction of a soil bacterial community. ISME J 9, 934–945 (2015).

25. G. Macedo, A. K. Olesen, L. Maccario, L. Hernandez Leal, P. v. d. Maas, D. Heederik, D. Mevius, S. J. Sørensen, H. Schmitt, Horizontal gene transfer of an IncP1 plasmid to soil bacterial community introduced by *Escherichia coli* through manure amendment in soil microcosms. Environ. Sci. Technol. 56, 11398–11408 (2022).

26. S. Musovic, G. Oregaard, N. Kroer, S. J. Sørensen, Cultivation-independent examination of horizontal transfer and host range of an IncP-1 plasmid among gram-positive and gram-negative bacteria indigenous to the barley rhizosphere. Appl. Environ. Microbiol. 72, 6687– 6692 (2006).

27. M. Arias-Andres, U. Klümper, K. Rojas-Jimenez, H.-P. Grossart, Microplastic pollution increases gene exchange in aquatic ecosystems. Environ. Pollut. 237, 253–261 (2018).

28. E. Wedel, C. Bernabe-Balas, M. Ares-Arroyo, N. Montero, A. Santos-Lopez, D. Mazel, B. Gonzalez-Zorn, Insertion sequences determine plasmid adaptation to new bacterial hosts. mBio 14, e03158–22 (2023).

29. H. E. Chia, E. N. G. Marsh, J. S. Biteen, Extending fluorescence microscopy into anaerobic environments. Curr. Opin. Chem. Biol. 51, 98–104 (2019).

30. M. C. Hansen, R. J. Palmer, C. Udsen, D. C. White, S. Molin, Assessment of GFP fluorescence in cells of *Streptococcus gordonii* under conditions of low pH and low oxygen concentration. Microbiology 147, 1383–1391 (2001).

31. H. Waseem, H. Saleem ur Rehman, J. Ali, M. J. Iqbal, M. I. Ali, “Chapter 14 - Global trends in ARGs measured by HT-qPCR platforms” in *Antibiotics and Antimicrobial Resistance Genes in the Environment*, M. Z. Hashmi, Ed. (Elsevier, 2020; https://www.sciencedirect.com/science/article/pii/B9780128188828000140)vol. 1 of *Advances in Environmental Pollution Research series*, pp. 206–222.

32. A. Gotz, K. Smalla, Manure Enhances Plasmid Mobilization and Survival of Pseudomonas putida Introduced into Field Soil. Appl Environ Microbiol 63, 1980–1986 (1997).

33. K. E. Hill, E. M. Top, Gene transfer in soil systems using microcosms. FEMS Microbiol. Ecol. 25, 319–329 (1998).

34. F. Berglund, T. Österlund, F. Boulund, N. P. Marathe, D. G. J. Larsson, E. Kristiansson, Identification and reconstruction of novel antibiotic resistance genes from metagenomes. Microbiome 7, 52 (2019).

35. K. J. Forsberg, A. Reyes, B. Wang, E. M. Selleck, M. O. A. Sommer, G. Dantas, The shared antibiotic resistome of soil bacteria and human pathogens. Science 337, 1107–1111 (2012).

36. S. D. Smith, J. Choi, N. Ricker, F. Yang, S. Hinsa-Leasure, M. L. Soupir, H. K. Allen, A. Howe, Diversity of Antibiotic Resistance genes and Transfer Elements-Quantitative Monitoring (DARTE-QM): a method for detection of antimicrobial resistance in environmental samples. Commun Biol 5, 1–9 (2022).

37. M. O. Press, A. H. Wiser, Z. N. Kronenberg, K. W. Langford, M. Shakya, C.-C. Lo, K. A. Mueller, S. T. Sullivan, P. S. G. Chain, I. Liachko, Hi-C deconvolution of a human gut microbiome yields high-quality draft genomes and reveals plasmid-genome interactions. bioRxiv, 198713 (2017).

38. C. W. Beitel, L. Froenicke, J. M. Lang, I. F. Korf, R. W. Michelmore, J. A. Eisen, A. E. Darling, Strain-and plasmid-level deconvolution of a synthetic metagenome by sequencing proximity ligation products. PeerJ 2, e415 (2014).

39. J. N. Burton, I. Liachko, M. J. Dunham, J. Shendure, Species-level deconvolution of metagenome assemblies with Hi-C-based contact probability maps. G3 (Bethesda) 4, 1339– 1346 (2014).

40. T. Stalder, M. O. Press, S. Sullivan, I. Liachko, E. M. Top, Linking the resistome and plasmidome to the microbiome. ISME J 13, 2437–2446 (2019).

41. E. Yaffe, D. A. Relman, Tracking microbial evolution in the human gut using Hi-C reveals extensive horizontal gene transfer, persistence and adaptation. Nat. Microbiol. 5, 343–353 (2020).

42. A. G. Kent, A. C. Vill, Q. Shi, M. J. Satlin, I. L. Brito, Widespread transfer of mobile antibiotic resistance genes within individual gut microbiomes revealed through bacterial Hi-C. Nat. Commun. 11, 4379 (2020).

43. L. Kalmar, S. Gupta, I. R. L. Kean, X. Ba, N. Hadjirin, E. M. Lay, S. P. W. de Vries, M. Bateman, H. Bartlet, J. Hernandez-Garcia, A. W. Tucker, O. Restif, M. P. Stevens, J. L. N. Wood, D. J. Maskell, A. J. Grant, M. A. Holmes, HAM-ART: An optimised culture-free Hi-C metagenomics pipeline for tracking antimicrobial resistance genes in complex microbial communities. PLOS Genet. 18, e1009776 (2022).

44. M. Marbouty, L. Baudry, A. Cournac, R. Koszul, Scaffolding bacterial genomes and probing host-virus interactions in gut microbiome by proximity ligation (chromosome capture) assay. Sci. Adv. 3, e1602105 (2017).

45. M. Marbouty, A. Cournac, J.-F. Flot, H. Marie-Nelly, J. Mozziconacci, R. Koszul, Metagenomic chromosome conformation capture (meta3C) unveils the diversity of chromosome organization in microorganisms. eLife 3, e03318 (2014).

46. M. Marbouty, A. Thierry, G. A. Millot, R. Koszul, MetaHiC phage-bacteria infection network reveals active cycling phages of the healthy human gut. eLife 10, e60608 (2021).

47. E. Top, M. Mergeay, D. Springael, W. Verstraete, Gene escape model: transfer of heavy metal resistance genes from Escherichia coli to Alcaligenes eutrophus on agar plates and in soil samples. Appl Environ Microbiol 56, 2471–2479 (1990).

48. J. W. Neilson, K. L. Josephson, I. L. Pepper, R. B. Arnold, G. D. Di Giovanni, N. A. Sinclair, Frequency of horizontal gene transfer of a large catabolic plasmid (pJP4) in soil. Appl Environ Microbiol 60, 4053–4058 (1994).

49. A. K. Lilley, J. C. Fry, M. J. Day, M. J. Bailey, *In situ* transfer of an exogenously isolated plasmid between *Pseudomonas spp.* in sugar beet rhizosphere. Microbiol 140, 27–33 (1994).

50. H. Heuer, K. Smalla, Plasmids foster diversification and adaptation of bacterial populations in soil. FEMS Microbiol. Rev. 36, 1083–1104 (2012).

51. S. Dealtry, G.-C. Ding, V. Weichelt, V. Dunon, A. Schlüter, M. C. Martini, M. F. D. Papa, A. Lagares, G. C. A. Amos, E. M. H. Wellington, W. H. Gaze, D. Sipkema, S. Sjöling, D. Springael, H. Heuer, J. D. van Elsas, C. Thomas, K. Smalla, Cultivation-independent screening revealed hot spots of IncP-1, IncP-7 and IncP-9 plasmid occurrence in different environmental habitats. PLOS ONE 9, e89922 (2014).

52. D. E. Wood, J. Lu, B. Langmead, Improved metagenomic analysis with Kraken 2. Genome Biology 20, 257 (2019).

53. R. I. Aminov, Horizontal gene exchange in environmental microbiota. Front Microbiol 2, 158 (2011).

54. V. Ivanova, E. Chernevskaya, P. Vasiluev, A. Ivanov, I. Tolstoganov, D. Shafranskaya, V. Ulyantsev, A. Korobeynikov, S. V. Razin, N. Beloborodova, S. V. Ulianov, A. Tyakht, Hi-C metagenomics in the ICU: Exploring clinically relevant features of gut microbiome in chronically critically ill patients. Front Microbiol 12 (2022).

55. V. F. Lanza, F. Baquero, J. L. Martínez, R. Ramos-Ruíz, B. González-Zorn, A. Andremont, A. Sánchez-Valenzuela, S. D. Ehrlich, S. Kennedy, E. Ruppé, W. van Schaik, R. J. Willems, F. de la Cruz, T. M. Coque, In-depth resistome analysis by targeted metagenomics. Microbiome 6, 11 (2018).

56. C. Gasc, P. Peyret, Revealing large metagenomic regions through long DNA fragment hybridization capture. Microbiome 5, 33 (2017).

57. A. K. Guitor, A. R. Raphenya, J. Klunk, M. Kuch, B. Alcock, M. G. Surette, A. G. McArthur, H. N. Poinar, G. D. Wright, Capturing the resistome: a targeted capture method to reveal antibiotic resistance determinants in metagenomes. Antimicrob. Agents Chemother. 64, e01324–19 (2019).

58. H. Heuer, C. T. T. Binh, S. Jechalke, C. Kopmann, U. Zimmerling, E. Krögerrecklenfort, T. Ledger, B. Gonzalez, E. Top, K. Smalla, IncP-1ε plasmids are important vectors of antibiotic resistance genes in agricultural systems: Diversification driven by class 1 integron gene cassettes. Front. Microbiol. 3, 2 (2012).

59. M. Popowska, A. Krawczyk-Balska, Broad-host-range IncP-1 plasmids and their resistance potential. Frontiers in Microbiology 4 (2013).

60. A. Jousset, C. Bienhold, A. Chatzinotas, L. Gallien, A. Gobet, V. Kurm, K. Küsel, M. C. Rillig, D. W. Rivett, J. F. Salles, M. G. A. van der Heijden, N. H. Youssef, X. Zhang, Z. Wei, W. H. G. Hol, Where less may be more: how the rare biosphere pulls ecosystems strings. ISME J 11, 853–862 (2017).

61. A. Reid, M. Buckley, The Rare Biosphere (American Society for Microbiology, Washington (DC), 2011; http://www.ncbi.nlm.nih.gov/books/NBK560451/)*American Academy of Microbiology Colloquia Reports*.

62. R. E. Fox, X. Zhong, S. M. Krone, E. M. Top, Spatial structure and nutrients promote invasion of IncP-1 plasmids in bacterial populations. ISME J 2, 1024–1039 (2008).

63. A. Schlüter, H. Heuer, R. Szczepanowski, L. J. Forney, C. M. Thomas, A. Pühler, E. M. Top, The 64 508 bp IncP-1β antibiotic multiresistance plasmid pB10 isolated from a waste-water treatment plant provides evidence for recombination between members of different branches of the IncP-1β group. Microbiol. 149, 3139–3153 (2003).

64. L. De Gelder, F. P. J. Vandecasteele, C. J. Brown, L. J. Forney, E. M. Top, Plasmid donor affects host range of promiscuous IncP-1 plasmid pB10 in an activated-sludge microbial community. Appl. Environ. Microbiol. 71, 5309–5317 (2005).

65. D. Hill, M. J. Morra, T. Stalder, S. Jechalke, E. Top, A. T. Pollard, I. Popova, Dairy manure as a potential source of crop nutrients and environmental contaminants. J. Environ. Sci. 100, 117–130 (2021).

66. C. M. Liu, M. Aziz, S. Kachur, P.-R. Hsueh, Y.-T. Huang, P. Keim, L. B. Price, BactQuant: An enhanced broad-coverage bacterial quantitative real-time PCR assay. BMC Microbiology 12, 56 (2012).

67. T. Větrovský, P. Baldrian, The variability of the 16S rRNA gene in bacterial genomes and its consequences for bacterial community analyses. PLoS One 8, e57923 (2013).

68. S. Chen, Y. Zhou, Y. Chen, J. Gu, fastp: an ultra-fast all-in-one FASTQ preprocessor. Bioinformatics 34, i884–i890 (2018).

69. H. Li, Aligning sequence reads, clone sequences and assembly contigs with BWA-MEM. arXiv:1303.3997 [q-bio] (2013).

70. W. Shen, H. Ren, TaxonKit: A practical and efficient NCBI taxonomy toolkit. JGG 48, 844– 850 (2021).

