## Supplementary File for "Detection of rare plasmid hosts using a targeted Hi-C approach"

### Supplementary Information

**Table S1: Soil microcosms with approximate relative abundances of each member[1]**

| **Soil Microcosm** | **Donor (D)** | **Recipient (R)** | **Transconjugant (T)** | **Soil Community** | **Transfer Frequency (T/R)** | **Hi-C** | **Hi-C +** |
|---|---|---|---|---|---|---|---|
| A | 4.6 x $10^{-5}$ | 0 | 5.2 x $10^{-3}$ | 0.95 | 1* | ✓ | [4] |
| B | 2.8 x $10^{-5}$ | 0.04 | 3.5 x $10^{-4}$ | 0.96 | $10^{-1}$ | ✓ | ✓ |
| C | 4.6 x $10^{-5}$ | 0.05 | 4.1 x $10^{-5}$ | 0.95 | $10^{-2}$ | ✓ | ✓ |
| D | 5.7 x $10^{-5}$ | 0.03 | 5.9 x $10^{-6}$ | 0.97 | $10^{-3}$ | ✓ | ✓ |
| E | 3.8 x $10^{-5}$ | 0.03 | 4.4 x $10^{-7}$ | 0.97 | $10^{-4}$ | ✓ | ✓ |
| F | 2.8 x $10^{-5}$ | 0.03 | 4.0 x $10^{-8}$ | 0.97 | $10^{-5}$ | [3] | ✓ |
| Control[2] | 4.7 x $10^{-5}$ | 0.03 | 0 | 0.97 | 0 | ✓ | ✓ |

[1] Donor: *E. coli* K12 MG1655Nal::gfp (pB10), Recipient: *P. putida* KT2442, Transconjugant: *P. putida* KT2442 (pB10)

[2] Control microcosm is soil inoculated with plasmid-free *E. coli* K12 MG1655 Nal::gfp and Recipient.

[3] Hi-C library for Soil Microcosm F was not sequenced. Preliminary results indicated this would be beyond the detection limit.

[4] Hi-C+ library for Soil Microcosm A was prepared, but not sequenced. Preliminary results indicated this would work well and we opted to leave it out in exchange for higher sequencing depth on the other Hi-C+ libraries.

[5] Soil bacteria were estimated using 16S rRNA quantification, see methods for more details. See Table S1 for representation of the cell densities of D, R and T as relative abundances, relative to the soil bacterial densities.

* The scenario in microcosm A mimics a case where all the recipients acquired the plasmid, making the transfer frequency 1

✓ Hi-C and Hi-C+ libraries were prepared and sequenced.

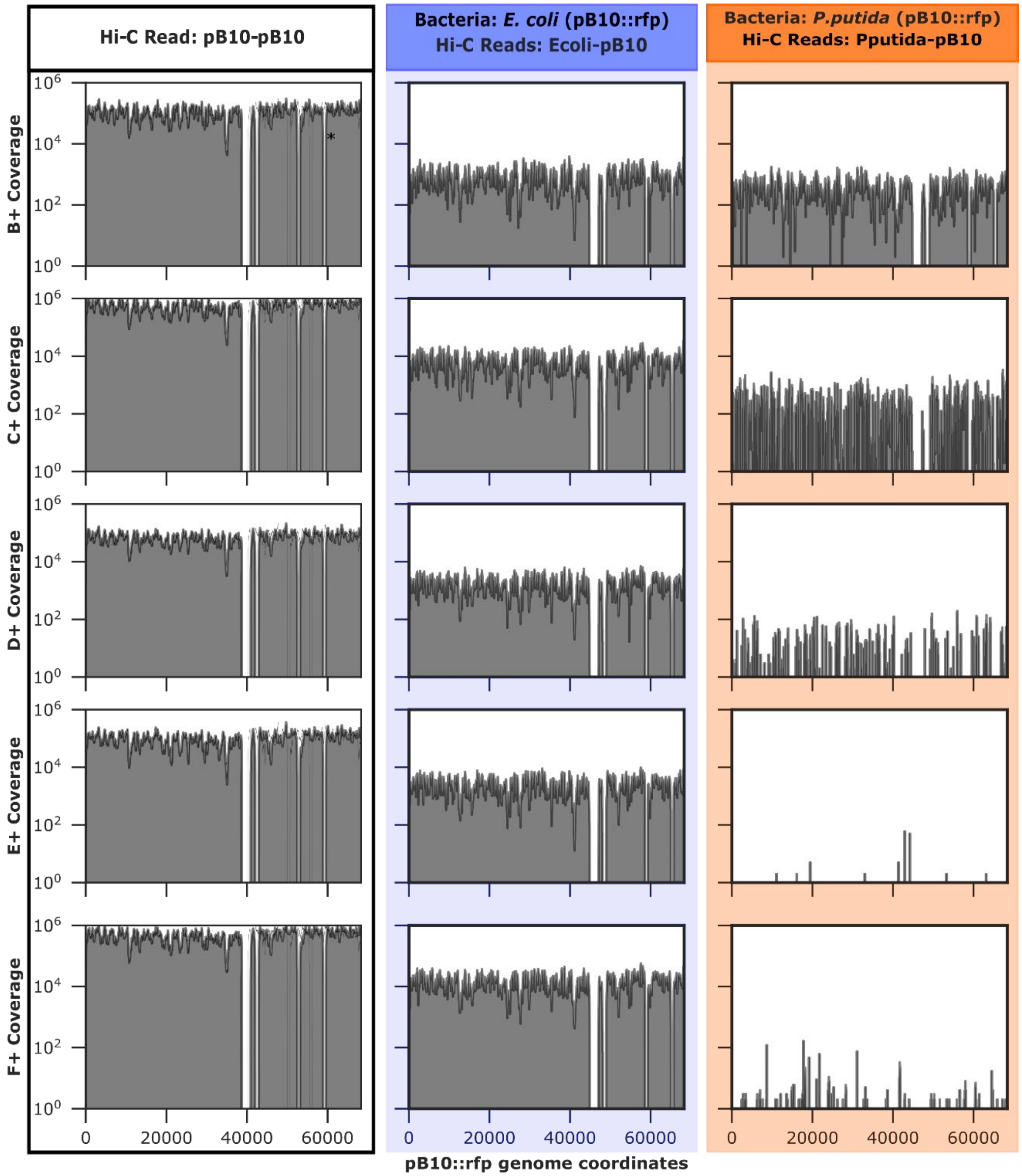


3 **Fig. S1: Coverage plots for all soil microcosms**. Each column is color-coded to indicate the Hi-
4 C+ reads for which data is presented.

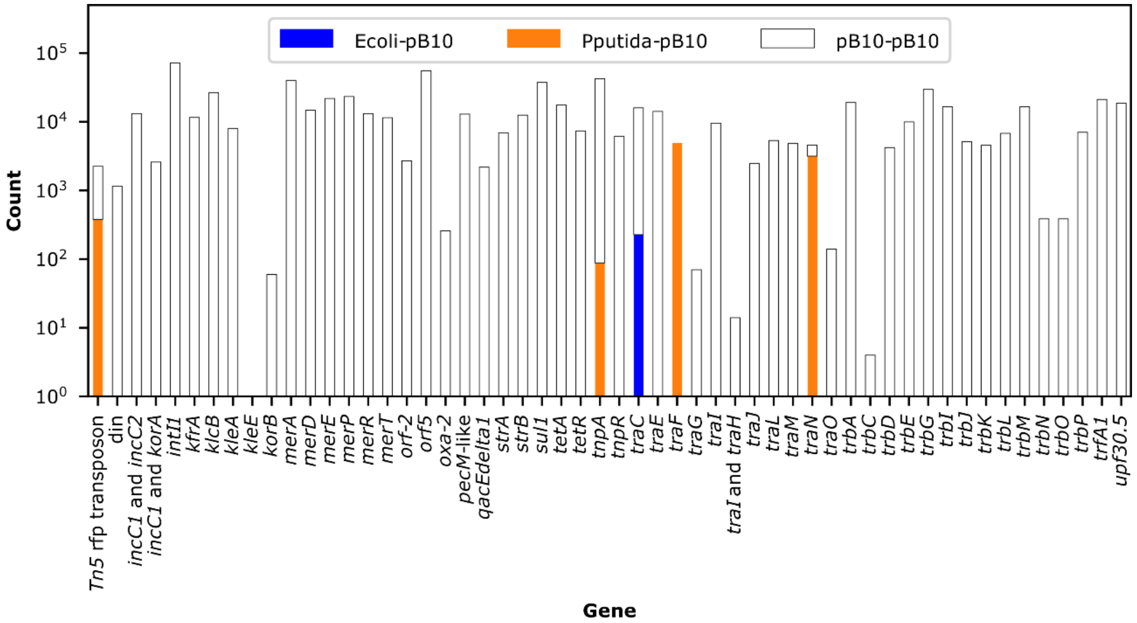


**Fig. S2. Plasmid genes detected in control soil microcosm.** Bars are color coded to indicate the number of each Hi-C read type that aligned to each gene.

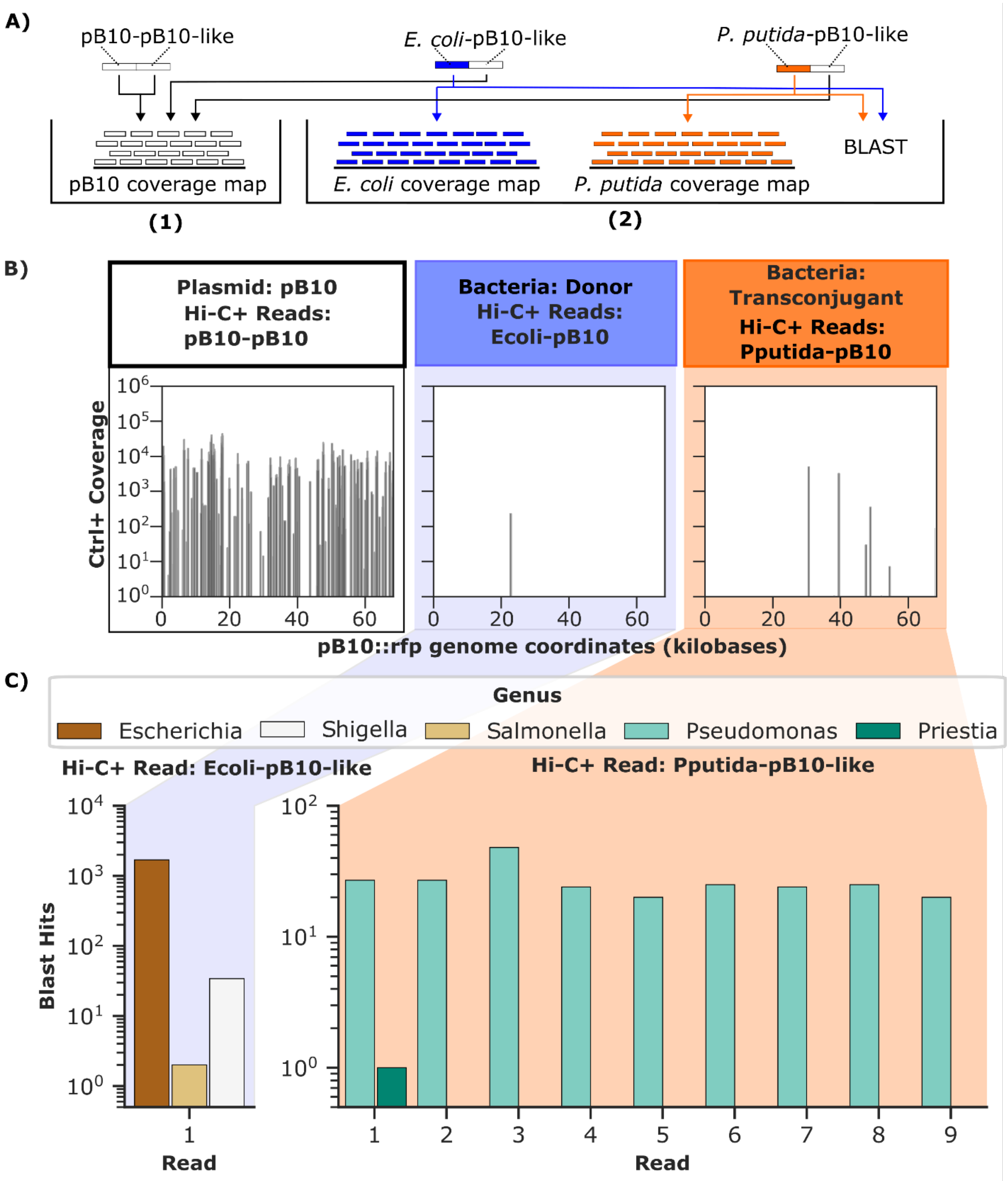


**Fig. S3. Investigation of plasmid-linked reads in control microcosm**. A) Visualization of the analysis carried out on the plasmid-linked reads in the control microcosm. Analysis 1 investigates where the pB10 segment of plasmid-like reads align on the plasmid reference genome. Analysis 2 determines where the *P. putida* KT2442 and *E. coli* MG1655 segments of plasmid-like reads align on their respective reference genomes. This is accompanied by a BLAST search of the sequence against the NCBI nucleotide database to determine whether the region to which the reads align is conserved across bacteria. B) Results from analysis 1, note these are also shown in Fig. 4. C) Results from analysis 2. Each unique read is numbered and shown on the x-axis, the number of reference sequences within each genus to which BLAST search mapped each read is shown on the y-axis. Only exact matches were included in the BLAST output.
